# Algebraic Morphogenesis Through Cochain Operators

**DOI:** 10.64898/2026.07.26.740851

**Authors:** Qichen Huang, Haoyang Guo

## Abstract

Cellular automata and graph reaction–diffusion systems encode local spatial interactions in different mathematical forms. We develop a cochain-operator calculus for these two settings. Over a finite field F_*q*_, every local rule on a finite neighborhood has a unique reduced polynomial representative. On an oriented line, the coboundary and endpoint maps recover the left and right shifts. Our main theorem shows that these operators, together with linear operations, constant cochains, and the degree-zero cup product, generate every finite-radius polynomial cellular automaton. Explicit formulas for Rules 30, 110, and 22 show how reflection-invariant linear coupling, directed transport, and nonlinear neighbor interactions enter the calculus. On a general graph, *d*^*^*d* is the unweighted combinatorial Laplacian and enters a graph reaction– diffusion recurrence. Over ℝ, the term − *Dd*^*^*d* with *D* ≥ 0 admits the usual diffusion interpretation; over F_*q*_, the corresponding expression defines modular coupling without an intrinsic order. In the morphogenetic examples, we therefore distinguish pattern-generating dynamics from finite-state observation and use the Betti numbers of active induced subcomplexes to summarize observed patterns. This yields a common algebraic representation without identifying real-valued diffusion with finite-field dynamics.

## 1. Wolfram’s rule as a polynomial

An elementary cellular automaton is a synchronous map on binary configurations *x*^*t*^ ∈ {0, 1} ^z^ determined by a local rule *f* : {0, 1}^3^ → {0, 1} . We begin with Wolfram’s Rule 30 in the standard numbering [19, 20]. It is a useful running example because its local rule already contains both linear and nonlinear operations.

Listing the 2^3^ neighborhoods in decreasing binary order, from 111 to 000, gives the transitions in Figure 1.

**FIGURE 1.**
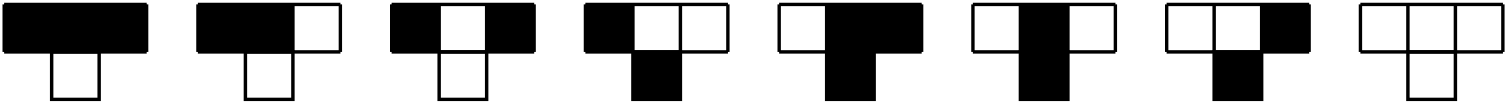
The local transition rules for Wolfram Rule 30.

**FIGURE 2.**
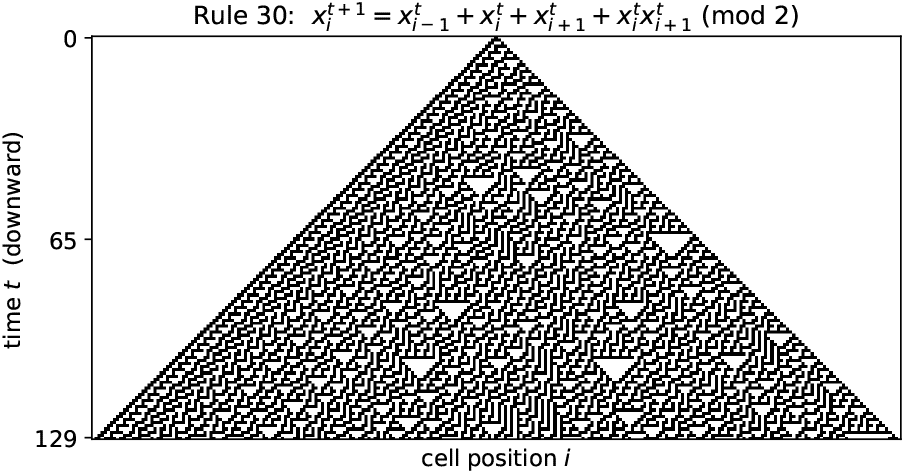
A finite portion of the Rule-30 orbit from a single active site, with time running downward. The orbit is obtained by iterating the polynomial (1.2) over F_2_.

The corresponding output word is 00011110_2_ = 30. Writing 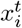 for the state at site *i* and time *t*, the update is

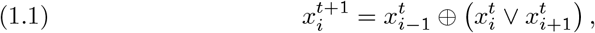

where ⊕ denotes XOR and ∨ denotes OR. After identifying 0, 1 with F_2_, these operations are represented by *A* ⊕ *B* = *A* + *B* and *A* ∨ *B* = *A* + *B* + *AB*. Thus (1.1) is the polynomial update

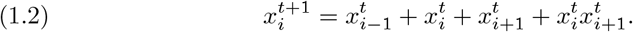

All operations in (1.2) are in F_2_.

### *Remark* 1.1.

Let Δ be the combinatorial Laplacian of the line. Over F_2_, (Δ*ϕ*)_*i*_ = *ϕ*_*i*−1_ + *ϕ*_*i*+1_, and hence

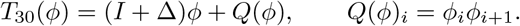

The affine operator *I* + Δ is Rule 150. The quadratic map *Q* is nonlinear and orientation-sensitive. It is not a pointwise reaction: it couples two adjacent sites. Section 3 treats diffusion together with pointwise polynomial reactions, while Section 4 shows how transported cup products generate *Q* and, more generally, every local polynomial interaction on the oriented line.

### 1.1. Relation to existing work

Polynomial descriptions of cellular automata are classical. Martin, Odlyzko and Wolfram [13] analyze additive rules through F_*p*_[*x*]*/*(*x*^*N*^ − 1), and Vivaldi [18] studies cellular-automaton evolution as a polynomial map over a finite field. Related algebraic methods have been developed for asynchronous updates [10] and for finite-state models of gene regulation and Boolean networks [9, 7, 17].

A separate literature uses cellular automata to model spatial pattern formation, including skin patterning and excitable media [22, 12]; broader accounts are given in [2, 1]. The present paper connects these two directions by placing polynomial local rules and the incidence Laplacian on the same cochain space. The formal result specific to this construction is Theorem 4.2: on an oriented line, endpoint maps and the cup product generate every finite-radius polynomial cellular automaton. The recurrence of Section 3 extends the notation to multispecies dynamics on a graph, while Section 5 distinguishes the algebraic finite-field model from real-valued diffusion.

## 2. Finite-field polynomial cellular automata

Three choices enter the passage from elementary cellular automata to finite spatial systems.

- **Spatial domain**. Sites may be the vertices of a finite graph rather than points of a regular lattice; edges record the pairs allowed to interact.
- **Neighborhood**. A local update may depend on any specified collection of *k* sites.
- **Alphabet**. The binary alphabet is replaced by a finite field F_*q*_, where *q* is a prime power. Its elements are algebraic state labels and carry no intrinsic order.

For a neighborhood state 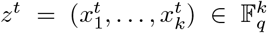, a local update is a map 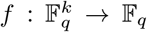. The following standard result identifies such maps with reduced polynomials.

### Theorem 2.1

(Polynomial normal form). *For every map* 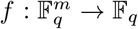 *there is a unique polynomial*

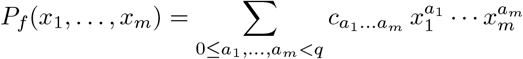

*with f*(*x*) = *P*_*f*_ (*x*) *for all* 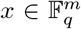. *Equivalently, evaluation is an isomorphism of* F_*q*_*-algebras*

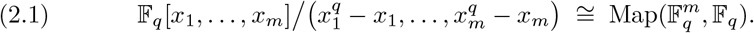

*Proof*. Reduction by 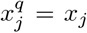 gives a representative of degree less than *q* in each variable, and both sides of (2.1) have dimension *q*^*m*^ over F_*q*_; it remains to show evaluation is injective. For *m* = 1, a polynomial of degree less than *q* vanishing at all *q* field elements is zero. For *m >* 1, regard a reduced polynomial as a polynomial in *x*_*m*_; fixing the first *m* − 1 coordinates, the one-variable case forces every coefficient to vanish, and induction on *m* finishes.

### Definition 2.2

Fix *r* ≥ 0. A *radius-r polynomial cellular automaton* on the line is specified by 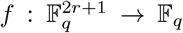 and acts by (*T*_*f*_ *ϕ*)_*i*_ = *f*(*ϕ*_*i*−*r*_, …, *ϕ*_*i*_, …, *ϕ*_*i*+*r*_). Its *affine part* is the sum of the constant and degree-one monomials of *P*_*f*_ ; the remaining monomials form its *nonlinear part*.

For *q* = 2 the relations 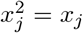 make *P*_*f*_ square-free, so a local rule is a unique sum of interaction monomials indexed by subsets of its neighborhood; Rule 30 has three singleton terms and one pair term, and the affine/nonlinear split of Definition 2.2 is exactly that of Remark 1.1.

### *Remark* 2.3.

The field assumption is used in Theorem 2.1. Over a general finite ring, not every function is polynomial, and the coordinate relations used in (2.1) do not provide the same unique reduced representative. Polynomial dynamics over finite rings remain possible, but an arbitrary local rule need not admit the normal form used here.

A prime-power alphabet can be taken to be 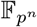 directly. Choosing a basis identifies this field with 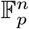 as a vector space, but the identification is not canonical and carries no biological interpretation by itself. More generally, a finite alphabet can be encoded as a subset of a sufficiently large field; such an encoding is a choice of labels and need not preserve a prescribed rule or any ordering of the states.

### *Remark* 2.4

(Coefficientwise lift). Reading a polynomial with coefficients 0, 1 over F_*p*_ for an odd prime *p* defines a useful new rule, but it is not a canonical scalar extension of the Boolean map: fields of different characteristic admit no unital homomorphism between them. We call this operation a *coefficientwise lift*. In equal characteristic, coefficients may instead be extended along the prime-subfield inclusion. Figure 4 and Appendix A show such lifts; the field elements carry no intrinsic order, and the colors displaying them are a reading aid, not part of the dynamics.

**FIGURE 3.**
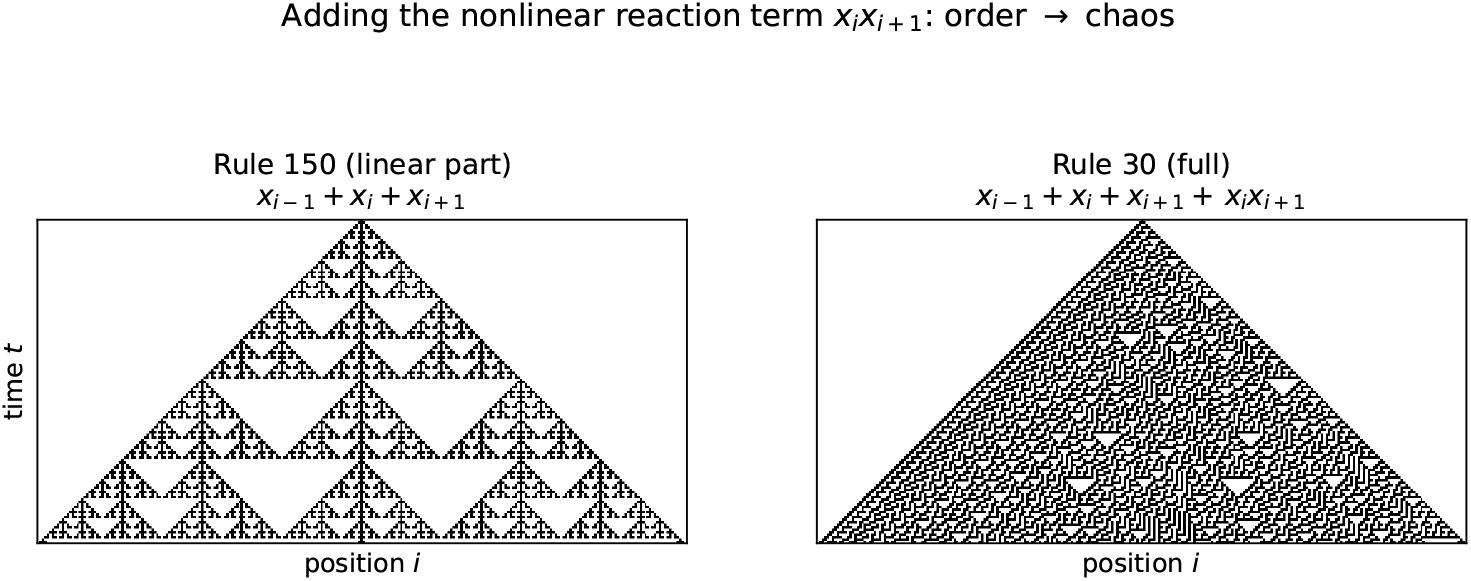
Orbits from the same single-site initial condition. *Left:* the affine rule *I* + Δ (Rule 150). *Right:* adding the orientation-sensitive quadratic term *Q*(*ϕ*)_*i*_ = *ϕ*_*i*_*ϕ*_*i*+1_ gives Rule 30.

**FIGURE 4.**
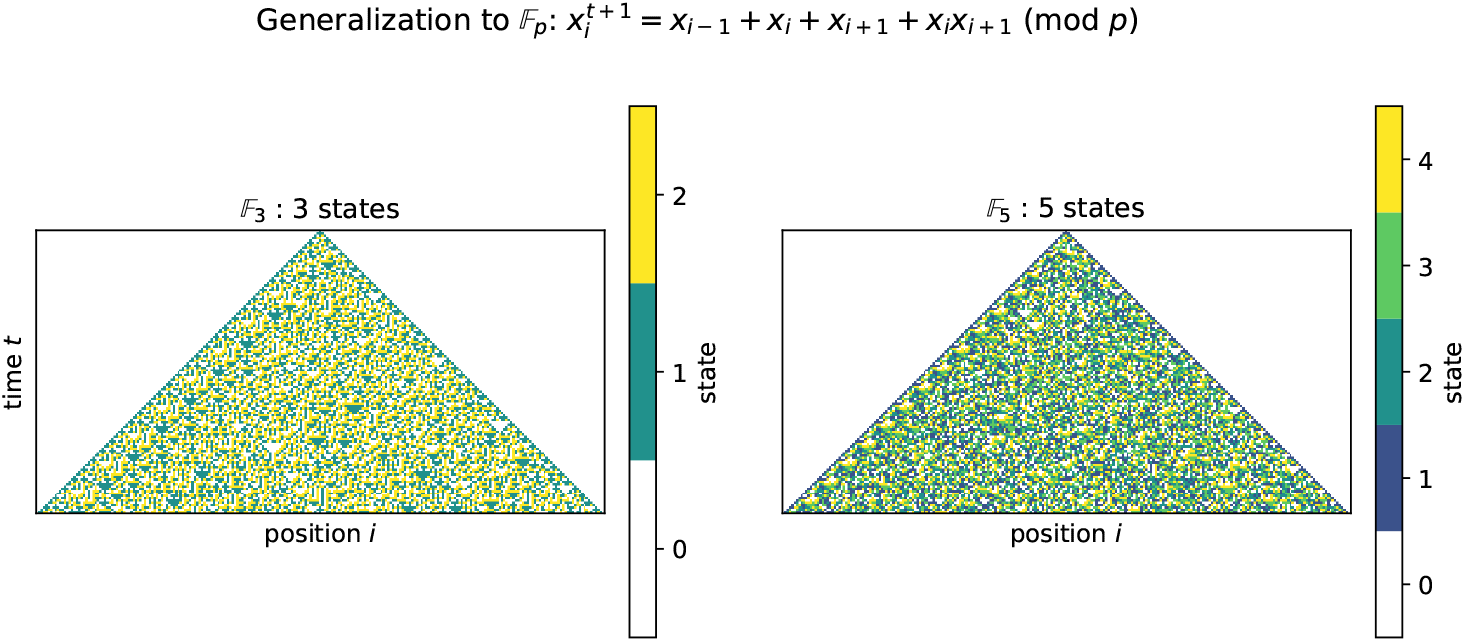
Coefficientwise lifts of Rule 30 (Remark 2.4). The polynomial *x*_*i*−1_ + *x*_*i*_ + *x*_*i*+1_ + *x*_*i*_*x*_*i*+1_ is iterated from one seed over F_3_ (left) and F_5_ (right); the state 0 is white. Appendix A gives further examples.

On a finite set of *N* sites, a specified local rule at each site determines a global map 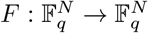. Each coordinate of *F* is a reduced polynomial in the states of the corresponding neighborhood. Translation invariance is an additional condition, not part of the polynomial representation itself.

### 2.1. Reflection symmetry on the line

For a radius-one rule, reflection interchanges the left and right variables: *ρ*(*ℓ, c, r*) = (*r, c, ℓ*). If *f* ◦ *ρ* = *f*, the induced global map commutes with reflection of the line. Otherwise the rule distinguishes the two orientations. This is the symmetry displayed in Figure 5 and characterized for affine Boolean rules in Proposition 3.4.

**FIGURE 5.**
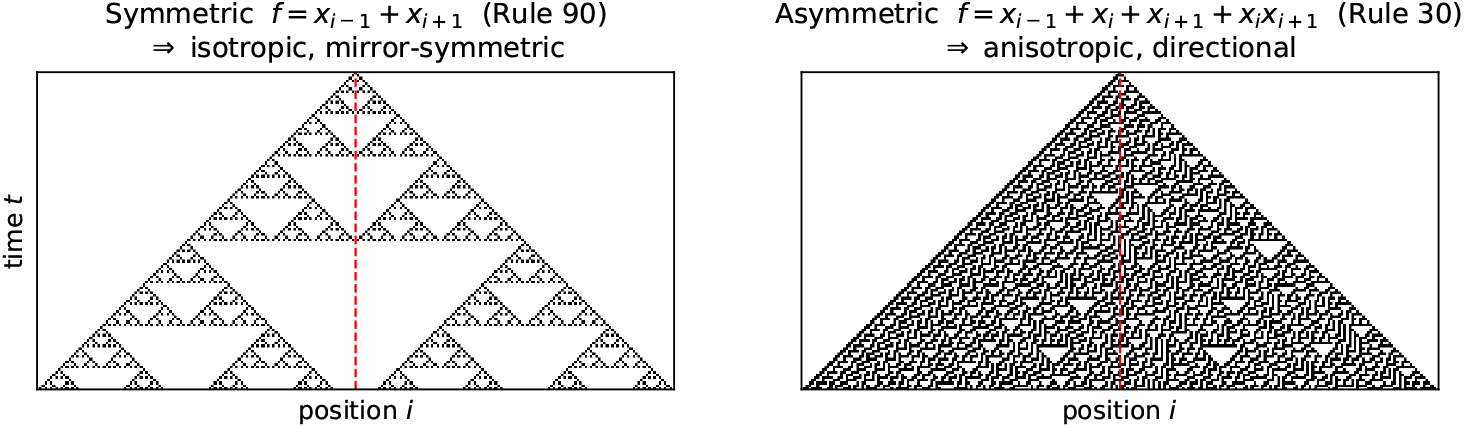
Reflection symmetry of two radius-one rules. *Left:* Rule 90, *f*(*ℓ, c, r*) = *ℓ* + *r*, is invariant under *ℓ* ↔ *r. Right:* the quadratic term *cr* in Rule 30 distinguishes the right neighbor and breaks reflection symmetry.

**FIGURE 6.**
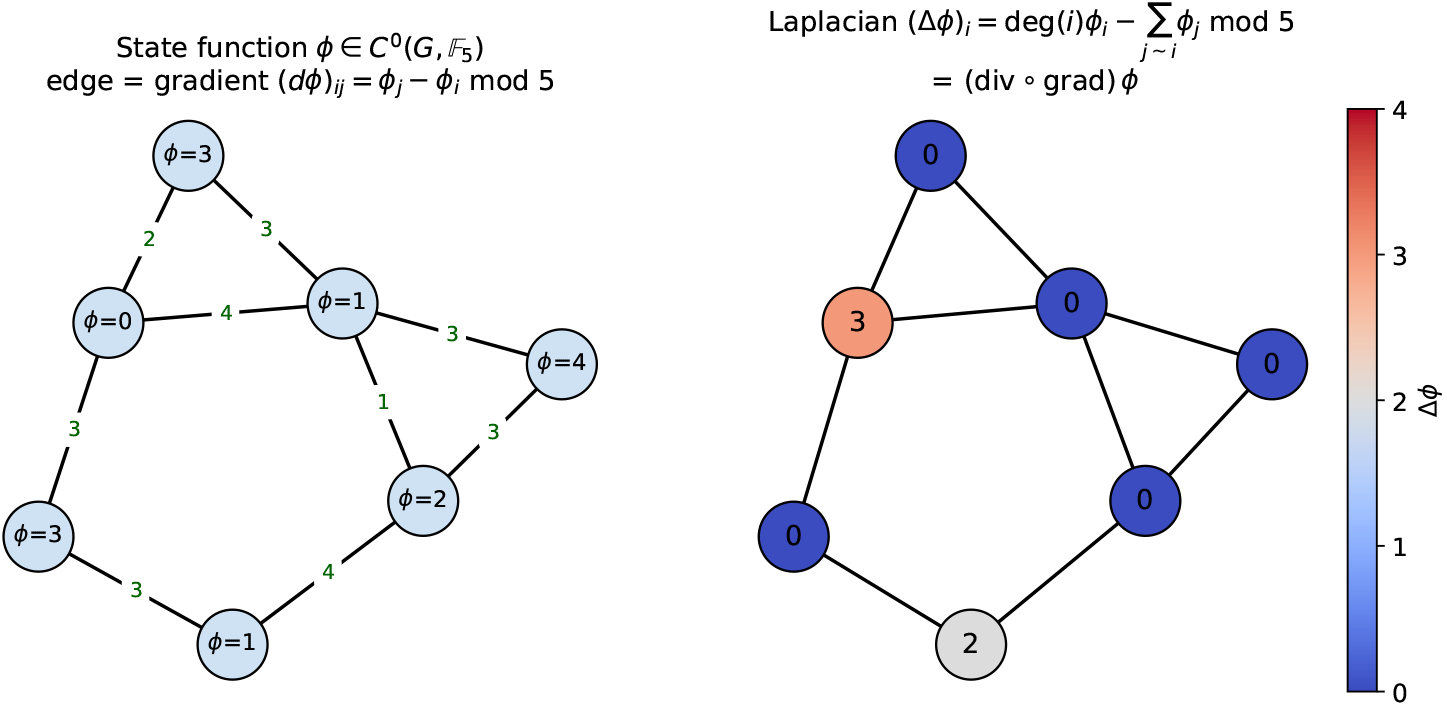
The cochain operators over F_5_. *Left:* a 0-cochain *ϕ* ∈ *C*^0^(*G*, F_5_) and the edge differences (*dϕ*)_*ij*_ = *ϕ*_*j*_ − *ϕ*_*i*_. *Right:* the graph Laplacian of (3.1). Reversing an edge changes the sign of its difference but not Δ (Proposition 3.2).

**FIGURE 7.**
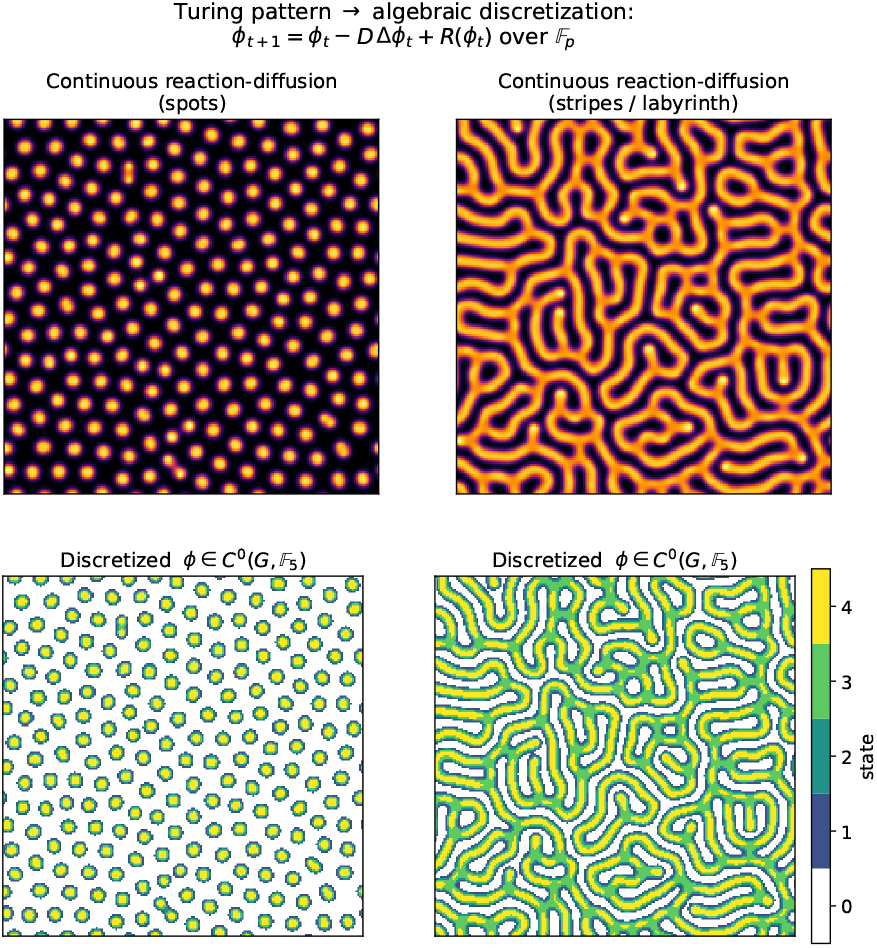
Real-valued Gray–Scott simulations and finite-field observations. *Top:* the two-component Gray–Scott system [5] with diffusion produces spot and labyrinthine patterns in two parameter regimes [15]. *Bottom:* each real-valued field is quantized into five labels and recorded as an F_5_-valued 0-cochain. The quantization is an observation step; no finite-field recurrence is used to generate these panels.

On an embedded graph, geometric isotropy is a stronger condition: it depends on the embedding, neighborhood geometry, and any edge weights, not only on permutation symmetry of the local polynomial. Proposition 3.7 gives the corresponding intrinsic statement in terms of graph automorphisms.

## 3. Cochain reaction–diffusion dynamics

Let **k** be either ℝ or a finite field. The incidence operators below are defined over either coefficient field, but their dynamical interpretation depends on that choice. Over ℝ, the term −*D*Δ with *D* ≥ 0 is the usual unweighted graph diffusion. Over F_*q*_, the same formula defines a linear coupling on a finite state space; it is not an averaging process because the field has no order or positivity. We use the same algebraic recurrence in both settings and reserve claims about physical diffusion and Turing instability for the real-valued model. Classical reaction–diffusion models are reviewed in [8]; the original pattern-forming mechanism is due to Turing [16].

### 3.1. The graph Laplacian over a field

Let *G* = (*V, E*) be a finite graph. We write vertex states as cochains in order to make the incidence structure explicit.

#### Definition 3.1

A *state* is a 0-cochain *ϕ* ∈ *C*^0^(*G*, **k**), that is, a map *ϕ* : *V* → **k**. An edge variable is a 1-cochain in *C*^1^(*G*, **k**); over ℝ it may be interpreted as a flux.

After choosing an ordering of *V*, the space *C*^0^(*G*, **k**) is identified with **k**^|*V* |^. The cochain notation retains the incidence map without requiring coordinates for the vertices. Fix an orientation on each edge *e* = (*u, v*). The coboundary *d* : *C*^0^ → *C*^1^ and its transpose *d*^*\**^ : *C*^1^ → *C*^0^, with respect to the standard coordinate pairings, are

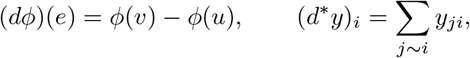

where the edge values are extended antisymmetrically, *y*_*ij*_ = −*y*_*ji*_. Their composite is the *combinatorial graph Laplacian* Δ = *d*^*\**^*d* : *C*^0^ → *C*^0^,

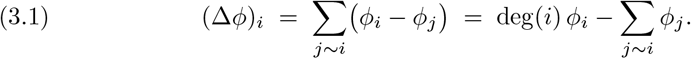

#### Proposition 3.2

*Over any field* **k**, *the operator* Δ = *d*^*\**^*d is independent of the chosen edge orientations and satisfies* (3.1) *together with* Δ**1** = 0, *where* **1** *is the constant cochain. If* **k** = ℝ *then*

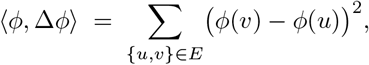

*so* ker Δ *consists of the functions constant on each connected component*.

*Proof*. Let *B* be the oriented incidence matrix, so *d* = *B* and Δ = *B*^T^*B*. Reversing an edge multiplies one row of *B* by − 1 and leaves *B*^T^*B* unchanged; expanding the product gives (3.1) and Δ**1** = 0. Over ℝ the displayed sum of squares vanishes exactly when the values agree across every edge.

Over a finite field, (3.1) is read in F_*q*_, with vertex degrees reduced modulo the characteristic. The sum-of-squares argument is then unavailable and Δ may have additional null vectors, although it remains well defined and orientation independent.

#### *Remark* 3.3

(Sign convention). The combinatorial Laplacian Δ = *d*^*\**^*d* = deg − *A* is the negative of the continuum operator: on a regular lattice of spacing *h*, − *h*^−2^Δ approximates ∇^2^. The real-valued diffusion term therefore has the sign −*D*Δ. The convention Δ = *d*^*\**^*d* also makes Δ positive semidefinite over ℝ.

#### 3.1.1. The discrete-time recurrence

Fix the cochain space *C*^0^(*G*, **k**) and let *ϕ*_*t*_ ∈ *C*^0^(*G*, **k**) for *t* ∈ N. Its forward difference is (*δϕ*)_*t*_ = *ϕ*_*t*+1_ − *ϕ*_*t*_. Given *D* ∈ **k** and a pointwise polynomial map *R* : *C*^0^(*G*, **k**) → *C*^0^(*G*, **k**), set

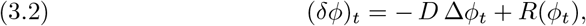

or equivalently

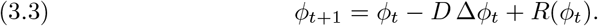

When **k** = ℝ, the diffusion interpretation assumes *D* ≥ 0 and a time step for which the chosen discretization is stable. When **k** = F_*q*_, all terms are evaluated in the field and (3.3) is a polynomial dynamical system.

##### Proposition 3.4

*Write a radius-one rule’s variables as* (*ℓ, c, r*) *and let reflection act by ρ*(*ℓ, c, r*) = (*r, c, ℓ*). *An affine Boolean rule f*(*ℓ, c, r*) = *αℓ* + *βc* + *γr* + *δ (α, β, γ, δ* ∈ F_2_*) is reflection invariant if and only if α* = *γ, in which case its global map is*

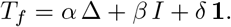

*Consequently the symmetric affine radius-one operators are exactly the* F_2_*-span of the identity I, the Laplacian* Δ *and the constant cochain* **1**.

*Proof*. Uniqueness in Theorem 2.1 gives *f*(*ℓ, c, r*) = *f*(*r, c, ℓ*) precisely when the coefficients of *ℓ* and *r* agree. Over F_2_, (Δ*ϕ*)_*i*_ = *ϕ*_*i*−1_ + *ϕ*_*i*+1_ (the diagonal 2*ϕ*_*i*_ vanishes), which yields *T*_*f*_ = *α*Δ + *βI* + *δ***1**. □

##### *Remark* 3.5

(Degree and reflection symmetry). Grading the algebraic normal form by degree and reflection symmetry separates the elementary rules into three classes.

- The eight **affine, reflection-invariant** rules form the F_2_-span of *I*, Δ, and **1**: Rules 0, 90, 150, 204 and their complements 255, 165, 105, 51.
- An **affine, reflection-breaking** rule cannot be written using *I*, Δ, and **1** alone. It requires oriented transport; Rules 170 = *ϕ*_*i*+1_ and 240 = *ϕ*_*i*_ − _1_ are the two shifts.
- A **nonlinear** rule contains monomials of degree at least two. Section 4 realizes these monomials as cup products of transported states. For Rule 30 the affine part is reflection invariant, while the term *ϕ*_*i*_*ϕ*_*i*+1_ breaks reflection symmetry.

Rule 110 combines the last two cases:

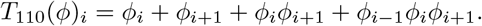

Its affine part breaks reflection symmetry, and its reduced polynomial also has quadratic and cubic terms. Over a general F_*q*_, the same classification uses the reduced polynomial of Theorem 2.1 in place of the Boolean algebraic normal form.

### 3.2. Two morphogens and the structure of the dynamics

Classical Turing instability is a diffusion-driven instability of a coupled system [16, 4]. For comparison, consider a scalar real-valued reaction–diffusion equation linearized at a homogeneous equilibrium *u*_*\**_. If *R*^*′*^(*u*_*\**_) *<* 0, then a Laplacian mode with eigen-value *λ* ≥ 0 has growth rate *R*^*′*^(*u*_*\**_) − *Dλ <* 0; ordinary diffusion cannot destabilize the equilibrium. At least two coupled components, or a mechanism outside this scalar setting, are therefore needed for a classical Turing instability.

For two components, take

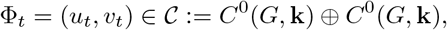

let ⅅ = diag(*D*_*u*_, *D*_*v*_), and let *R* = (*R*_*u*_, *R*_*v*_) act pointwise. Applying Δ to each component gives

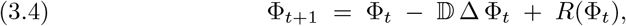

that is, componentwise,

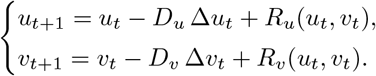

In the real-valued Turing setting one takes positive, typically unequal, diffusion coefficients. Over F_*q*_, the coefficients are field elements and (3.4) instead defines a finite polynomial dynamical system. The same formula extends to any number *s* of components and has the following two structural properties.

#### Proposition 3.6

*Over* F_*q*_, *the recurrence* (3.4) *is a polynomial dynamical system on* 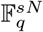 *(s the number of species, N* = | *V* |*). Every orbit is eventually periodic, and the sum of its transient length and period is at most* 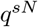.

*Proof*. Δ is linear and *R* is polynomial, so the update is a polynomial self-map of a set with 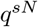 elements. Among the first 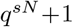 states two coincide, and determinism makes the subsequent orbit periodic. □

#### Proposition 3.7

*Let σ* ∈ Aut(*G*) *act on cochains by pullback. If the diffusion coefficients are the same at every vertex and the reaction R acts pointwise, by the same polynomial at each vertex, the update map T of* (3.4) *is equivariant: T* (*σ*^*\**^Φ) = *σ*^*\**^*T* (Φ).

*Proof*. Graph automorphisms preserve degrees and adjacency, hence commute with Δ by (3.1); pointwise evaluation of a fixed *R* commutes with pullback. □

Thus Δ does not break any automorphism already present in the graph. Symmetry breaking in an individual trajectory may still arise from its initial condition or from an instability. Any explicit directional preference in the update, however, must enter through the graph, spatially varying coefficients, or oriented neighbor data such as those of Remark 4.3.

#### 3.2.1. Source-driven gradients

In the real-valued one-component model, take the linear reaction *R*(*ϕ*) = − *kϕ* with *k >* 0, let *D >* 0, and impose a boundary source of strength *C*_0_. On the continuum half-line, the steady problem

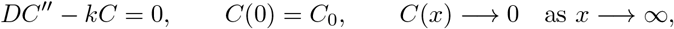

has the solution *C*(*x*) = *C*_0_*e*^−*x/λ*^ with 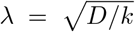. Thresholding this profile partitions the domain into positional regions, as in Wolpert’s French-flag model of positional information [21].

This source-driven gradient is distinct from a diffusion-driven instability. In the simulations of Figure 8, random initial fluctuations without a source decay toward a homogeneous state. Classical spontaneous pattern selection requires additional coupled nonlinear feedback, as in the two-component system above.

**FIGURE 8.**
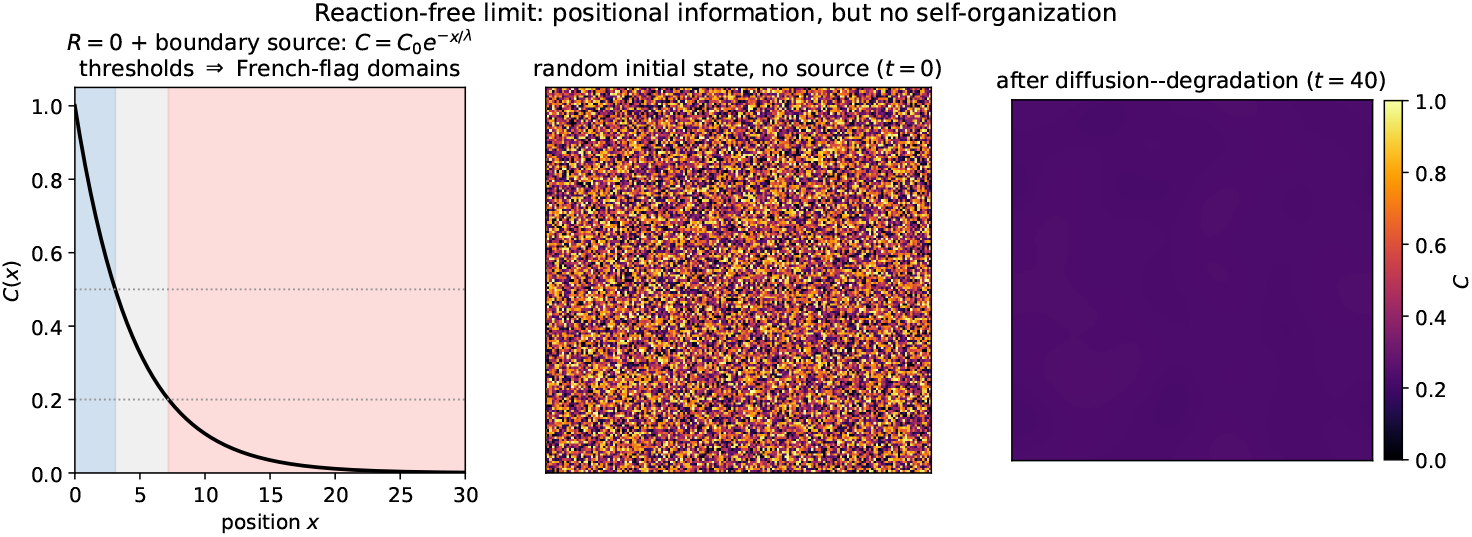
Diffusion–degradation with the linear reaction *R*(*ϕ*) = − *kϕ. Left:* a boundary source gives *C* = *C*_0_*e*^−*x/λ*^; two thresholds divide the profile into three positional regions. *Middle and right:* without a source, random initial fluctuations decay toward a homogeneous state.

**FIGURE 9.**
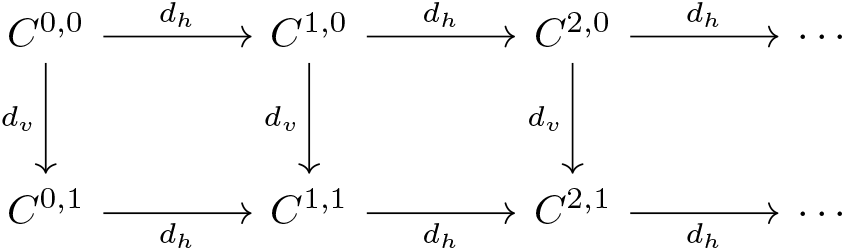
The product bicomplex *C*^*p,q*^ = *C*^*p*^(*X*; **k**) ⊗ *C*^*q*^(*I*_*T*_ ; **k**). The horizontal differential is the spatial coboundary; the vertical differential is the signed temporal coboundary.

### 3.3. The space–time bicomplex and pattern complexes

Let *X* be a finite ordered simplicial complex with 1-skeleton *G*, and let *I*_*T*_ be the simplicial interval with vertices 0, …, *T* and oriented edges [*t, t* + 1]. For a coefficient field **k**, set

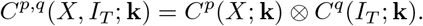

The horizontal differential is *d*_*h*_ = *d*_*X*_ ⊗ 1. On *C*^*p,q*^ define the signed vertical differential by *d*_*v*_ = (−1)^*p*^1 ⊗ *d*_*I*_. Then

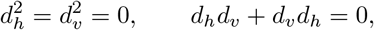

so *C*^•,•^(*X, I*_*T*_ ; **k**) is a bicomplex. Since *I*_*T*_ is one-dimensional, only the rows *q* = 0, 1 are nonzero.

A trajectory (*ϕ*_0_, …, *ϕ*_*T*_ ) is a *C*^0^(*X*; **k**)-valued 0-cochain on the vertices of *I*_*T*_, hence an element of *C*^0,0^. Its temporal coboundary satisfies

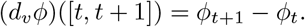

The recurrence (3.3) can therefore be imposed edgewise as

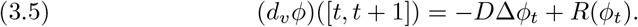

The bicomplex comes from the product *X × I*_*T*_ ; equation (3.5) is a generally non-linear constraint on a trajectory, not a bicomplex differential.

The cohomology of the fixed complex *X* describes the topology of the spatial domain, not the topology of a state on that domain. For a binary state *ϕ* ∈ *C*^0^(*X*; F_2_), define the active induced subcomplex

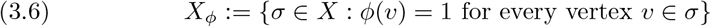

and the pattern Betti numbers

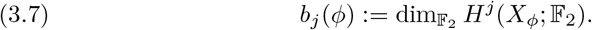

Thus *b*_0_ counts active connected components, while *b*_1_ counts active cycles not filled by active 2-simplices. For a multistate field one must first choose which labels are active; because F_*q*_ has no intrinsic order, this choice is part of the observation map. The complexes 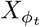 need not be nested as *t* varies, so (3.7) is a time-slice observable; tracking classes through time would require an additional persistence construction.

## 4. The cochain algebra and the generation theorem

The recurrence (3.4) still treats *R* as an external polynomial: Δ = *d*^*\**^*d* has a meaning on the complex, while *R* is specified separately. Cochains also carry a product, the cup product [6], which represents pointwise polynomial reactions. Directed neighbor interactions require additional incidence data—the endpoint maps introduced below—and diffusion uses the adjoint *d*^*\**^. We therefore keep these ingredients explicit rather than attributing all three operations to the differential graded algebra alone.

### 4.1. The cochain algebra

Fixing the vertex order of *X*, the cup product ⌣: *C*^*a*^ *× C*^*b*^ → *C*^*a*+*b*^ is

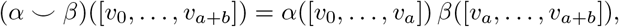

making (*C*^•^(*X*, **k**), *d*, ⌣) a differential graded algebra: it obeys the Leibniz rule *d*(*α* ⌣ *β*) = *dα* ⌣ *β* + ( − 1)^|*α*|^*α* ⌣ *dβ* and descends to the cup product on cohomology. In degree zero it is the pointwise product (*ϕ* ⌣ *ψ*)(*i*) = *ϕ*_*i*_*ψ*_*i*_; the mixed product *C*^0^ *× C*^1^ → *C*^1^ evaluates the *tail* of each edge, (*ϕ* ⌣ *ψ*)(*e*) = *ϕ*(tail *e*) *ψ*(*e*). This tail/head asymmetry reflects the chosen vertex order. To turn it into a directed update on vertices, one must also specify which incident edge is read at each vertex.

### 4.2. Transports

On the oriented line, with *e*_*i*_ = [*i, i* + 1], the endpoint contractions *∂*^+^, *∂*^−^ : *C*^1^ → *C*^0^ read off the outgoing and incoming edge at *i, ∂*^+^*ψ*(*i*) = *ψ*(*e*_*i*_) and *∂*^−^*ψ*(*i*) = *ψ*(*e*_*i* − 1_) (so that *d*^*\**^ = *∂*^−^ − *∂*^+^). They remember which incident edge is right and which is left, and recover the shifts.

#### Lemma 4.1

*On the oriented line the right and left shifts on C*^0^ *satisfy*

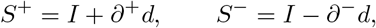

*that is, S*^+^*ϕ*(*i*) = *ϕ*_*i*+1_ *and S*^−^*ϕ*(*i*) = *ϕ*_*i*−1_.

*Proof*. At vertex *i*, (*I* + *∂*^+^*d*)*ϕ* = *ϕ*_*i*_ + (*ϕ*_*i*+1_ − *ϕ*_*i*_) = *ϕ*_*i*+1_, while (*I* − *∂*^−^*d*)*ϕ* = *ϕ*_*i*_ − (*ϕ*_*i*_ − *ϕ*_*i*−1_) = *ϕ*_*i*−1_. □

Every neighbor value is thus a {*d, ∂*^*±*^}-word applied to *ϕ*, and every monomial of a local rule is a cup product of transported copies of *ϕ*.

### 4.3. The generation theorem

#### Theorem 4.2

(Generation). *Let* 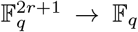 *be a radius-r local rule with reduced polynomial P*_*f*_ . *Its global cellular automaton on the line is*

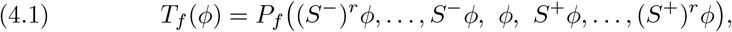

*where polynomial multiplication is the degree-zero cup product. Consequently every local polynomial cellular automaton is generated by addition, scalar multiplication, constant cochains, d, ∂*^*±*^, *and* ⌣. *Conversely, every expression formed from finitely many shifts, constant cochains, linear combinations, and degree-zero cup products defines a translation-equivariant polynomial cellular automaton of finite radius. For a radius-one rule, the reflection-invariant sum of the two neighbor terms is a multiple of* Δ = *d*^*\**^*d over* F_2_ *and lies in the span of I and* Δ *over a general* F_*q*_, *since S*^−^*ϕ* + *S*^+^*ϕ* = 2*ϕ* − Δ*ϕ; a central linear term contributes separately to I. Higher-order terms are sums of cup products of transported copies. Thus the full local update is represented in the operator calculus generated by linear operations, constants, d, ∂*^*±*^, *and* ⌣.

*Proof*. Evaluating (4.1) at vertex *i* substitutes *ϕ*_*i*_ − _*r*_, …, *ϕ*_*i*+*r*_ into *P*_*f*_, hence gives (*T*_*f*_ *ϕ*)_*i*_ = *f*(*ϕ*_*i*_ − _*r*_, …, *ϕ*_*i*+*r*_); Lemma 4.1 expresses every shift using *d* and *∂*^*±*^. Conversely, a finite expression contains only finitely many shifts, and its evaluation at *i* is a polynomial in the corresponding finite neighborhood, independent of *i*. □

### 4.4. Worked examples

Over F_2_, using *∂*^+^(*ϕ* ⌣ *dϕ*)_*i*_ = *ϕ*_*i*_(*ϕ*_*i*+1_ − *ϕ*_*i*_) = *ϕ*_*i*_*ϕ*_*i*+1_ + *ϕ*_*i*_, Rules 30 and 110 can be written as

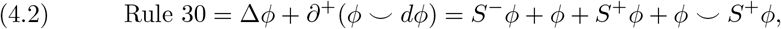

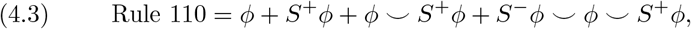

and the symmetric but nonlinear reference rule is

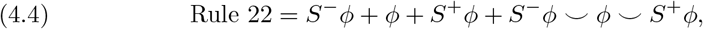

with pointwise polynomials *P*_110_(*ℓ, c, r*) = *c* + *r* + *cr* + *ℓcr* and *P*_22_(*ℓ, c, r*) = *ℓ* + *c* + *r* + *ℓcr*. In (4.2) the nonlinear term *∂*^+^(*ϕ ⌣ dϕ*) formally resembles the one-sided product *ϕ ∂*_*x*_*ϕ*, but over F_2_ this notation does not by itself define a continuum or Burgers limit. Its orientation dependence (Remark 3.5) comes from the outgoing endpoint map *∂*^+^ together with the tail convention of ⌣. Equation (4.3) displays both features noted in Remark 3.5: its affine part *ϕ* + *S*^+^*ϕ* is asymmetric, and its remaining terms are quadratic and cubic transported products. The construction is not special to F_2_: over F_*q*_ (Figure 4) the Rule-30 polynomial is *ϕ*+*S*^−^*ϕ*+*S*^+^*ϕ*+*ϕ ⌣ S*^+^*ϕ*. Over fields of odd characteristic the diagonal 2*ϕ*_*i*_ of Δ no longer vanishes, so the two shifts are a more direct expression of the linear part than Δ alone. In every characteristic, the update remains a word in *d, ∂*^*±*^, and ⌣, together with linear operations and constants.

#### Remark 4.3

(Ports). The shifts in Theorem 4.2 use the two labeled ports of the oriented line. On a regular graph with neighbor maps *σ*_*j*_ : *V* → *V* the same proof uses *S*_*j*_*ϕ* = *ϕ* ◦ *σ*_*j*_; without such port data an unlabeled graph canonically supports only rules invariant under permutations of indistinguishable neighbors, the Laplacian Δ being of this type (Proposition 3.7).

### 4.5. Pointwise and transported products

The generation theorem places the recurrence (3.4) and local cellular automata in a common operator calculus. A *pointwise* reaction, such as the activator–inhibitor coupling *R*(*u, v*) of a Turing system, is represented by linear combinations and cup products of same-vertex species cochains; for example, *u ⌣ v* carries no transport. A *cellular-automaton interaction*, such as Rule 30’s *ϕ ⌣ S*^+^*ϕ*, is a cup product of *transported* copies of a single species. These operations use related but distinct pieces of structure: ⌣ for pointwise multiplication, *d*^*\**^*d* for the unweighted graph Laplacian, and *d* together with *∂*^*±*^ for directed transport. A transported product is no longer pointwise, and its equivariance is therefore relative to the port data of Remark 4.3; without those data Proposition 3.7 applies only to neighbor-permutation-invariant constructions.

## 5. Representation, analysis, and the generation gap

The construction provides a common notation, not an equivalence of dynamics. The differential and cup product can be defined over either ℝ or a finite field, while the adjoint, the port maps, and any rounding or thresholding operation supply additional structure. In particular, the interpretation of a formula depends on its coefficient system.

### *Remark* 5.1.

With the standard cochain inner products, Δ = *d*^*\**^*d* in (3.1) is the unweighted combinatorial graph Laplacian. It is defined on an irregular graph without choosing lattice coordinates, but it is not automatically an exact discretization of the continuum Laplace operator. A geometry-sensitive PDE discretization may instead require weights or a discrete Hodge star. Over ℝ, the unweighted operator nevertheless gives a well-defined graph reaction–diffusion model; Figure 11 records one numerical example on a synthetic irregular graph.

Over F_*p*_, the same written expression defines a modular cellular automaton rather than physical diffusion. A finite field has no order compatible with its field operations, so its addition cannot be interpreted intrinsically as averaging values from high concentration to low. In the experiment of Figure 10, the particular modular update tested does not smooth a random field in the way its real-valued counterpart does. This is a statement about that update and discretization, not a claim that finite-field cellular automata cannot form spatial patterns.

**FIGURE 10.**
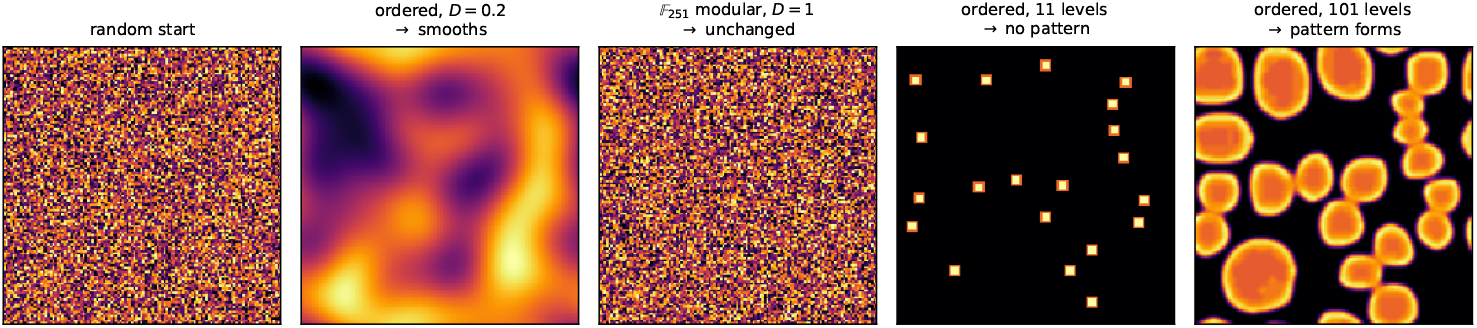
Two numerical probes of the role of the coefficient system (Remark 5.1). *Panels 1–3:* the real and modular updates are applied to the same random field. The real-valued graph diffusion smooths this field, whereas the modular update used here does not. *Panels 4–5:* the Gray–Scott update is followed by quantization to two different ordered resolutions. The lower-resolution run loses the pattern visible at the higher resolution. These panels illustrate the displayed parameter choices rather than a general threshold in field size or resolution.

Ordered quantization gives a different model again: the state has finitely many ordered levels, but rounding is an external operation rather than finite-field arithmetic. The examples in Figure 10 show that the outcome of one Gray–Scott discretization depends strongly on the chosen resolution; they do not establish a universal minimum number of levels. The threshold construction in Appendix B follows a second route. Its state and output are binary, but the short- and long-range neighborhood averages are compared as ordered real numbers before thresholding. Thus it is a binary-state threshold automaton, not a computation carried out entirely inside F_2_. Its output can nonetheless be read directly as a cochain *ϕ* ∈ *C*^0^(*X*, F_2_). In both routes the same complex may support the generative dynamics and the topological observation, but the coefficient system and the map between the two layers are part of the model.

Concretely, the cup product runs over ℝ as readily as over F_*p*_. Taking the Gray– Scott reaction [5, 15] *R*_*u*_(*u, v*) = − *uv*^2^ + *F* (1 − *u*), *R*_*v*_(*u, v*) = *uv*^2^ − (*F* + *κ*)*v* on an irregular Delaunay graph, with all products the degree-zero cup product and a clip Π(*z*) = min*{*1, max*{*0, *z}}*, gives the real-valued recurrence

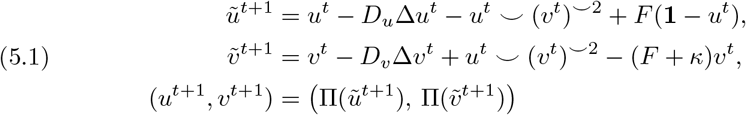

for which three computed regimes are shown in Figure 11. The clipping operation uses the order on ℝ and is not part of the cochain algebra. Passage to the analysis layer is likewise an explicit *observation map*: for a nonnegative real cochain *z* and *ε >* 0, first set

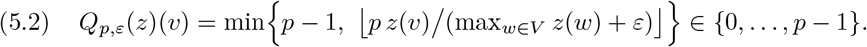

**FIGURE 11.**
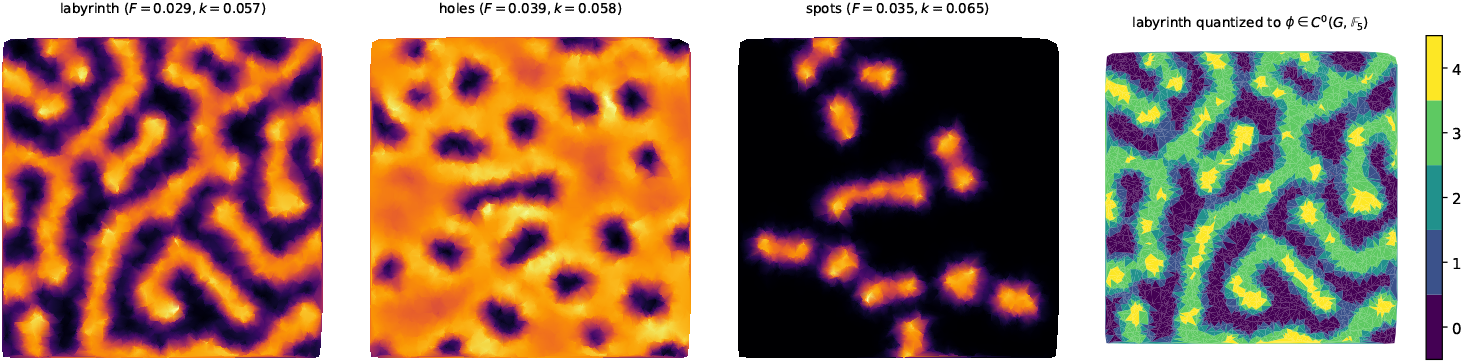
Computed states of the real-valued recurrence (5.1), with Δ = *d*^*\**^*d*, on a synthetic irregular Delaunay graph with 5000 vertices. The three parameter choices produce labyrinth-, hole-, and spot-like states in this simulation. The rightmost panel applies the observation map (5.2) with *p* = 5 to the labyrinth state, recording the resulting labels as *ϕ* ∈ *C*^0^(*G*, F_5_).

Identifying each integer label with its residue class gives a cochain in *C*^0^(*X*, F_*p*_). The map uses the order on ℝ; it is not a field homomorphism or a conjugacy of dynamics.

## 6. Biological examples

The ocellated lizard *Timon lepidus* provides a biological example close to the two-layer picture of Remark 5.1. Its adult dorsal pattern is organized on a quasi-hexagonal lattice of scales whose green and black states evolve as a probabilistic cellular automaton. Manukyan et al. derived this mesoscopic automaton from a continuous Turing reaction–diffusion model, with the geometry of the scales separating microscopic and mesoscopic length scales [11]. Subsequent three-dimensional simulations on reconstructed skin geometries showed that variation in skin thickness can by itself produce scale-by-scale coloration and automaton-like switching in the model [3].

For the present formalism, one may take the scales as vertices of their dual adjacency graph *G* and encode the two observed colors by a cochain in *C*^0^(*G*, F_2_). This is a mathematical coarse-graining of the biological system; it should not be confused with the particular quantizer in (5.2). Zebrafish stripes provide a related, but not identical, example. Experiments on melanophores and xanthophores found opposing short- and long-range effects, and the interaction network inferred from those experiments satisfies a local-activation, long-range-inhibition condition used in Turing models [14]. This motivates the threshold construction in Appendix B, but does not identify that construction with the cellular dynamics of the fish.

## 7. Conclusion

We developed a cochain-operator calculus that separates three ingredients of local spatial dynamics: symmetric linear coupling, directed transport, and nonlinear interaction. On an oriented line, the coboundary and endpoint maps recover the left and right shifts, while degree-zero cup products encode polynomial interactions among transported states. Together with linear operations and constant cochains, these operators generate every finite-radius polynomial cellular automaton over a finite field (Theorem 4.2). On a general graph, the incidence coboundary and its transpose supply the combinatorial Laplacian *d*^*\**^*d*. Polynomial cellular automata and graph reaction–diffusion recurrences can therefore be expressed in a common operator language, while port-dependent transport remains distinct from intrinsic graph diffusion.

The coefficient system is part of the model, not merely a change of notation. Over ℝ, the term − *Dd*^*\**^*d*, with *D* ≥ 0, has the usual graph-diffusion interpretation; over a finite field, the corresponding expression defines modular coupling without an intrinsic order. Quantization, clipping, and threshold comparison must therefore be specified as observation or update operations; they are not finite-field analogues of physical diffusion. This distinction permits real-valued and discrete pattern generators to be compared without identifying their dynamics.

For a binary state, the cohomology of its active induced subcomplex provides a time-slice summary: *b*_0_ counts active connected components, while *b*_1_ counts active cycles not filled by active 2-simplices. These invariants are deliberately coarse. Natural next steps are to study their stability under the observation map, organize thresholded states into multiscale filtrations, and track classes across the nonnested sequence 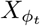.

## Appendix A. Cellular automata over finite fields

To illustrate the trichotomy of Remark 3.5, we iterate the same 0–1 polynomial for each elementary rule over F_*q*_, for *q* = 2, 3, 5, 7; when *q >* 2, this is the coefficien-twise lift of Remark 2.4. Figure 12 collects the three nonlinear rules of that remark, each grown from a single seed. The algebraic form is unchanged across each row, while the finite spacetime plots change visibly with the field.

**FIGURE 12.**
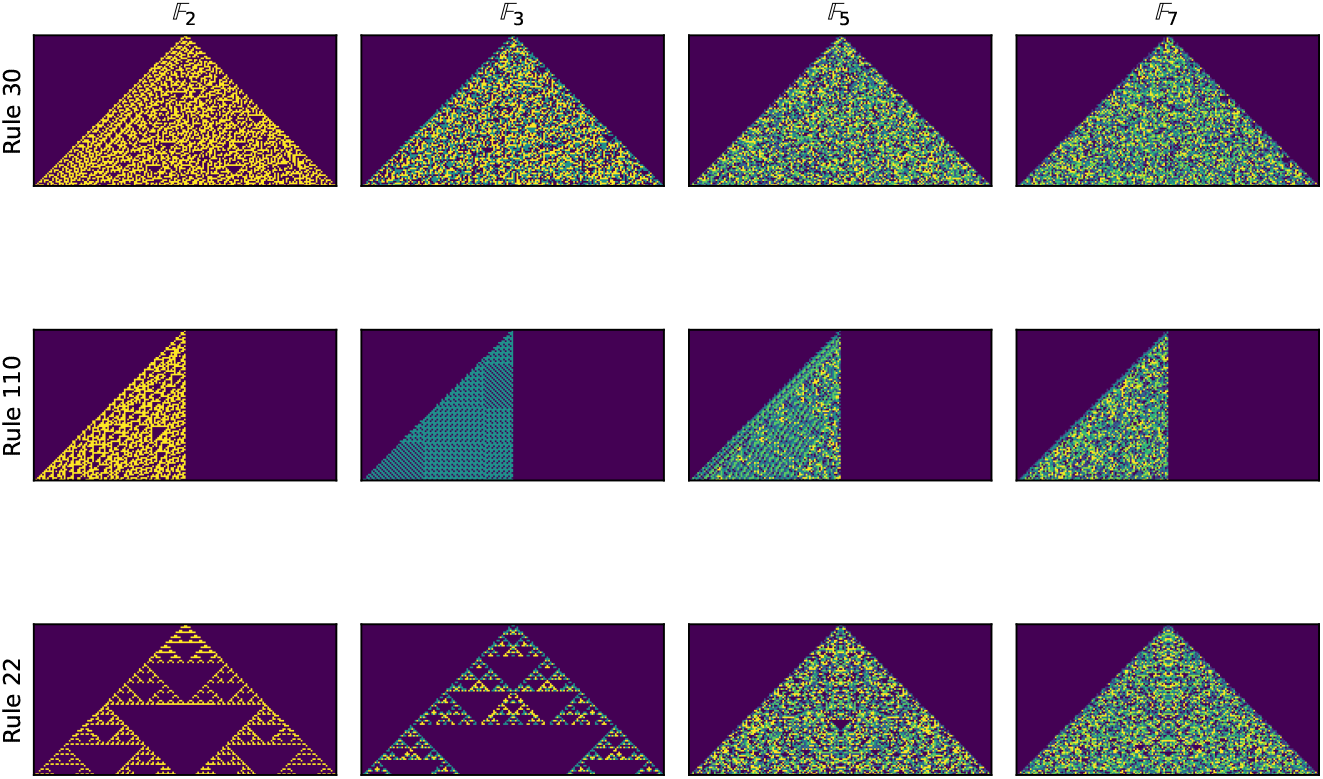
The nonlinear elementary rules 30, 110, and 22 iterated from a single active seed over F_*q*_, for *q* = 2, 3, 5, 7, using the same 0–1 polynomial across each row (time downward, with the quiescent state 0 dark). Rule 30 is directed through its quadratic term, Rule 110 already has a directed affine part, and Rule 22 is the reflection-symmetric nonlinear reference. The colors distinguish field elements but do not impose an order on them.

## Appendix B. A binary-state threshold generator and its cohomology

The threshold generator of Remark 5.1 extends from a lattice to a finite graph by replacing Euclidean disks with short- and long-range graph neighborhoods. At each update it compares the corresponding normalized active fractions and then applies a pointwise threshold, producing a binary state *ϕ* ∈ *C*^0^(*G*, F_2_). The comparison uses ordered real arithmetic, although the state before and after each update is binary. The output can be read by its F_2_ Betti numbers *b*_0_ = dim *H*^0^ (connected components) and *b*_1_ = dim *H*^1^ (enclosed loops), computed on the 2-complex *X* so that *b*_1_ counts pattern-scale holes rather than mesh triangles (§3.3). Figure 13 shows the three regimes on a Delaunay graph of 5000 vertices. In this run their Betti-number pairs are distinct: the spot-like state has many components and no holes, the labyrinthine state has both, and the hole-like state has relatively few components and many holes.

**FIGURE 13.**
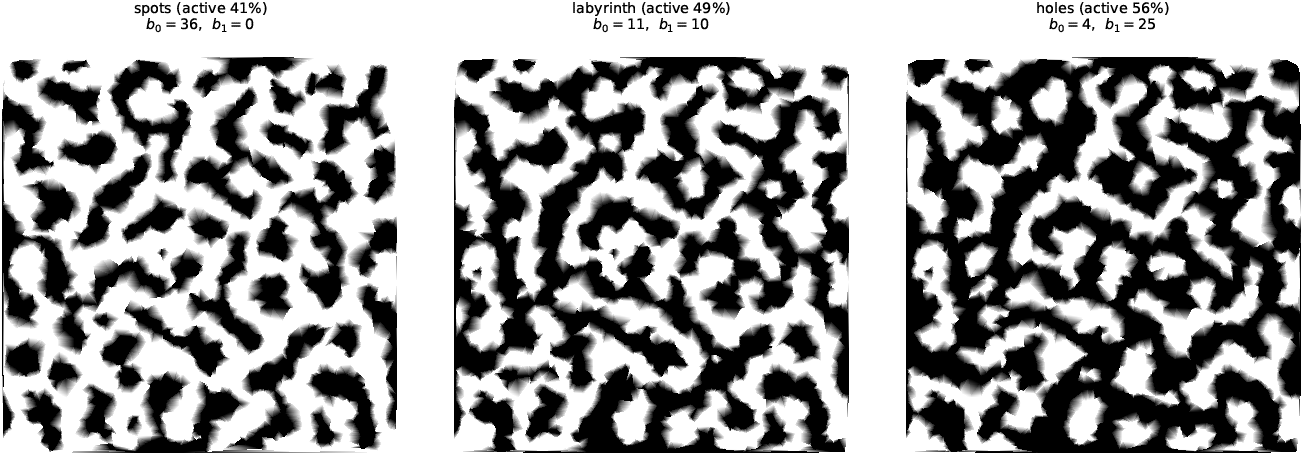
A binary-state threshold generator on an irregular Delaunay graph of 5000 vertices (active vertices black), with three outputs obtained by varying the inhibition weight. Each output is recorded as a 0-cochain *ϕ* ∈ *C*^0^(*G*, F_2_); its F_2_ Betti numbers *b*_0_, *b*_1_, computed on the induced subcomplex of the filled Delaunay complex *X*, are printed above the corresponding panel.

## References

[1] A. Deutsch and S. Dormann, Cellular Automaton Modeling of Biological Pattern Formation, 2nd ed., Birkhäuser, Boston, MA, 2017.

[2] G. B. Ermentrout and L. Edelstein-Keshet, Cellular automata approaches to biological modeling, J. Theoret. Biol. 160 (1993), no. 1, 97–133.

[3] A. Fofonjka and M. C. Milinkovitch, Reaction-diffusion in a growing 3D domain of skin scales generates a discrete cellular automaton, Nat. Commun. 12 (2021), Paper No. 2433.

[4] A. Gierer and H. Meinhardt, A theory of biological pattern formation, Kybernetik 12 (1972), no. 1, 30–39.

[5] P. Gray and S. K. Scott, Autocatalytic reactions in the isothermal, continuous stirred tank reactor: oscillations and instabilities in the system A + 2B → 3B; B → C, Chem. Eng. Sci. 39 (1984), no. 6, 1087–1097.

[6] A. Hatcher, Algebraic Topology, Cambridge Univ. Press, Cambridge, 2002.

[7] A. S. Jarrah, R. Laubenbacher, B. Stigler, and M. Stillman, Reverse-engineering of polynomial dynamical systems, Adv. in Appl. Math. 39 (2007), no. 4, 477–489.

[8] S. Kondo and T. Miura, Reaction-diffusion model as a framework for understanding biological pattern formation, Science 329 (2010), no. 5999, 1616–1620.

[9] R. Laubenbacher and B. Stigler, A computational algebra approach to the reverse engineering of gene regulatory networks, J. Theoret. Biol. 229 (2004), no. 4, 523–537.

[10] M. Macauley, J. McCammond, and H. S. Mortveit, Dynamics groups of asynchronous cellular automata, J. Algebraic Combin. 33 (2011), no. 1, 11–35. arXiv:0808.1238.

[11] L. Manukyan, S. A. Montandon, A. Fofonjka, S. Smirnov, and M. C. Milinkovitch, A living mesoscopic cellular automaton made of skin scales, Nature 544 (2017), no. 7649, 173–179.

[12] M. Markus and B. Hess, Isotropic cellular automaton for modelling excitable media, Nature 347 (1990), no. 6288, 56–58.

[13] O. Martin, A. M. Odlyzko, and S. Wolfram, Algebraic properties of cellular automata, Comm. Math. Phys. 93 (1984), no. 2, 219–258.

[14] A. Nakamasu, G. Takahashi, A. Kanbe, and S. Kondo, Interactions between zebrafish pigment cells responsible for the generation of Turing patterns, Proc. Natl. Acad. Sci. USA 106 (2009), no. 21, 8429–8434.

[15] J. E. Pearson, Complex patterns in a simple system, Science 261 (1993), no. 5118, 189–192.

[16] A. M. Turing, The chemical basis of morphogenesis, Philos. Trans. Roy. Soc. London Ser. B 237 (1952), no. 641, 37–72.

[17] A. Veliz-Cuba, A. S. Jarrah, and R. Laubenbacher, Polynomial algebra of discrete models in systems biology, Bioinformatics 26 (2010), no. 13, 1637–1643.

[18] F. Vivaldi, Cellular automata and finite fields, Phys. D 79 (1994), no. 2–4, 115–131.

[19] S. Wolfram, Statistical mechanics of cellular automata, Rev. Modern Phys. 55 (1983), no. 3, 601–644.

[20] S. Wolfram, A New Kind of Science, Wolfram Media, Champaign, IL, 2002.

[21] L. Wolpert, Positional information and the spatial pattern of cellular differentiation, J. Theoret. Biol. 25 (1969), no. 1, 1–47.

[22] D. A. Young, A local activator-inhibitor model of vertebrate skin patterns, Math. Biosci. 72 (1984), no. 1, 51–58.

